# Expression of a membrane-targeted fluorescent reporter disrupts auditory hair cell mechanoelectrical transduction and causes profound deafness

**DOI:** 10.1101/2020.09.18.303743

**Authors:** Angela Ballesteros, Tracy S. Fitzgerald, Kenton J. Swartz

**Affiliations:** Molecular Physiology and Biophysics Section. National Institute of Neurological Disorders and Stroke. National Institutes of Health. Bethesda, MD. 20892; Mouse Auditory Testing Core. National Institute on Deafness and other Communication Disorders. National Institutes of Health. Bethesda, MD. 20892

**Author notes:** Correspondence to: Angela Ballesteros, Kenton J. Swartz.

**Keywords:** Deafness, fluorescent reporter mice, membrane-targeted reporter, hair cell, prestin, mechanotransduction

## Abstract

The reporter mT/mG mice expressing a membrane-targeted fluorescent protein are becoming widely used to study the auditory and vestibular system due to its versatility. Here we show that high expression levels of the fluorescent mtdTomato reporter affect the function of the sensory hair cells and the auditory performance of mT/mG transgenic mice. Auditory brainstem responses and distortion product otoacoustic emissions revealed that adult mT/mG homozygous mice are profoundly deaf, whereas heterozygous mice present high frequency loss. We explore whether this line would be useful for studying and visualizing the membrane of auditory hair cells by airyscan super-resolution confocal microscopy. Membrane localization of the reporter was observed in hair cells of the cochlea, facilitating imaging of both cell bodies and stereocilia bundles without altering cellular architecture or the expression of the integral membrane motor protein prestin. Remarkably, hair cells from mT/mG homozygous mice failed to uptake the FM1-43 dye and to locate TMC1 at the stereocilia, indicating defective mechanoelectrical transduction machinery. Our work emphasizes that precautions must be considered when working with reporter mice and highlights the potential role of the cellular membrane in maintaining functional hair cells and ensuring proper hearing.

## INTRODUCTION

Transgenic mice expressing reporter genes are valuable tools for studying a wide range of cellular processes and for identifying specific tissues, cell populations and subcellular structures (Abe et al., 2013; Li et al., 2018; Stewart et al., 2009; Vacaru et al., 2014). Reporter mouse strains encoding fluorescent proteins allow easy visualization of target genes over endogenous genes (Abe et al., 2013; Vacaru et al., 2014), as long as the reporter is expressed in sufficient quantities and is stable. Directed reporter mouse lines are preferred over random genomic insertion to minimize side effects and avoid extreme expression levels of the transgene, which may lead to infertile or not viable mice [1].

Fluorescent fusion proteins that localize to the plasma membrane provide information on cell morphology and membrane dynamics in living cells (Abe et al., 2013; Larina et al., 2009; Shioi et al., 2011; Watanabe et al., 2019). The mT/mG mice (strain No. 007676, The Jackson laboratories. B6.129(Cg)-Gt(ROSA)26Sortm4(ACTB-tdTomato,-EGFP) express a cell membrane-localized tandem dimer of the Tomato (mtdTomato) fluorescent protein in all cells and tissues (Muzumdar et al., 2007). The mtdTomato cassette is inserted in the Rosa26 locus of C57BL/6J mice, a locus widely used to directly insert reporter genes since it supports the ubiquitous transcriptional activity of transgenes without affecting the animal viability or fertility (Li et al., 2018; Zambrowicz et al., 1997). The inserted cassette includes a chimeric cytomegalovirus enhancer/chicken β-actin promoter (pCA) that allows for strong and persistent expression of mtdTomato (Madisen et al., 2010; Tchorz et al., 2012). In addition, two loxP sites flanking the mtdTomato transgene and its polyadenylation (pA) signal allow the removal of the fluorescent protein after Cre recombination, and subsequent expression of membrane-targeted GFP (mGFP) cloned downstream of mtdTomato (Muzumdar et al., 2007)(Figure 1A). This design makes the mT/mG mice a versatile tool to visualize the cell membrane, to identify a specific cell type, and to track cell linages, because the double transgenic offspring of crosses between mT/mG mice and cellspecific Cre mice express mtdTomato in non-Cre recombined cells and mGFP in Cre-recombined cells.

**Figure 1:**
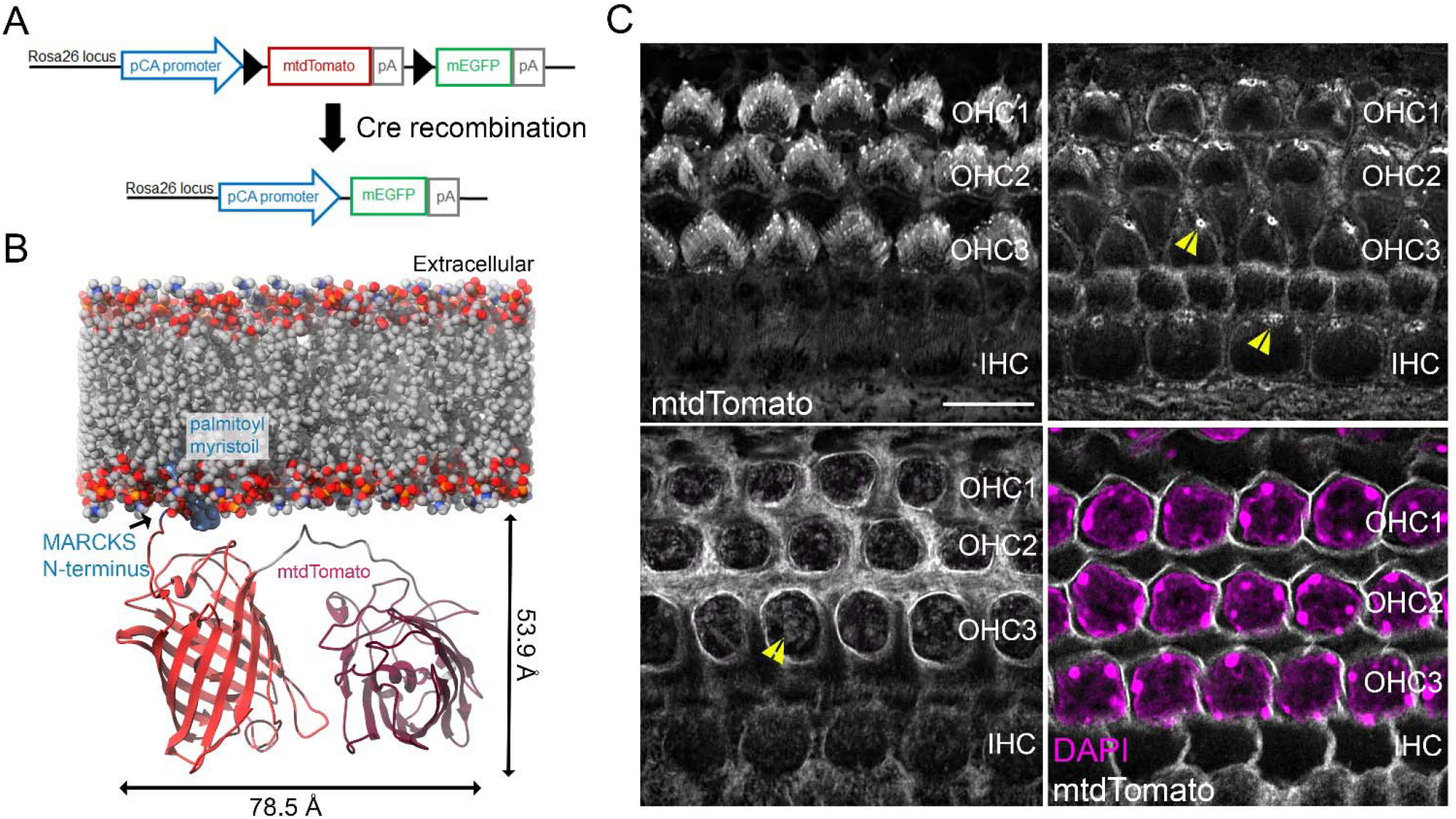
The mT/mG reporter mice. **A)** Schematic representation of the genomic design of the mT/mG reporter strain. The transgene inserted in the Rosa26 genetic locus includes a chimeric CMV/ β-actin (pCA) promoter that drives the expression of a membrane-targeted tandem dimer Tomato (mtdTomato) fluorescent protein. The mtdTomato reporter is located between two LoxP sites (black triangles) so that after Cre-mediated recombination, mtdTomato is excised, and a membrane-targeted green fluorescent protein (mEGFP) is expressed. **B)** Cartoon representation of a mtdTomato protein in a lipid bilayer. A structural model of the tdTomato was modified with a palmitoyl and myristoyl group in its MARCKS terminus (blue) and embedded into a phosphatidylcholine membrane to visualize the dimensions of the reporter. **C)** mtdTomato labeling of the inner (IHC) and outer (OHC) auditory hair cells membrane at the stereocilia (top left), cuticular plate (top right), subcuticular plate (bottom left) and the cell membrane at the nucleus (bottom right) region. DAPI (magenta) was included in the mounting media to label the cell nucleus. Scale bar represents 10 μm.

Hearing impairment is the most prevalent sensory defect in humans, with around 466 million people worldwide presenting disabling hearing loss. The mT/mG mice are a valuable tool to study and visualize several tissues and cells (Hasegawa et al., 2015; Liu et al., 2013; Muzumdar et al., 2007; Portal et al., 2019), and they are becoming widely used to study both the auditory and vestibular systems. In vestibular hair cells, mT/mG mice have been used for lineage tracing of hair cells of the cristae (Slowik et al., 2013) and of supporting cells in the murine utricle (Burns et al., 2012) and saccule (Jiang et al., 2017), and to track exosome release from supporting and hair cells of the murine utricle (Breglio et al., 2020). Likewise, to study auditory hair cells, mT/mG mice have been crossed with several specific Cre strains to label and differentiate supporting cells of the murine cochlea (Korrapati et al., 2013), for lineage tracing of cochlear hair cells (Mizutari et al., 2013; Walters et al., 2015), and as a reporter of the Cre activity in murine hair cells (Landin Malt et al., 2019). mtdTomato has also been used to visualize the morphology of the organ of Corti (Puria et al., 2011; Soons et al., 2015) and to study the lateral diffusion of membrane proteins in murine outer hair cells (OHCs) (Yamashita et al., 2015). However, despite the increasing implementation of mT/mG reporter mice in the auditory field, a thorough characterization of the auditory phenotype of mT/mG mice is still lacking.

In the present study, we investigated whether the mT/mG mice would be useful for studying the auditory hair cell membrane using super resolution confocal microscopy. Although the expression of the mtdTomato reporter greatly facilitates imaging of the hair cell body and stereocilia bundle without altering the cellular architecture, high levels of expression of the reporter impair the function of sensory hair cells and the auditory performance of mT/mG transgenic mice. Auditory brainstem response (ABR) and disortion product otoacoustic emissions (DPOAEs) recordings revealed that adult mT/mG homozygous mice (mT/mG^Tg/Tg^) are profoundly deaf, whereas heterozygous mice (mT/mG^Tg/+^) present high frequency hearing loss when compared to littermates lacking the transgene (mT/mG^+/+^). We provide some insights into the mechanism underlying this auditory phenotype of mT/mG mice. In particular, while the overall morphology of the auditory hair cells and expression and localization of the protein motor prestin was preserved, auditory hair cells from mT/mG^Tg/Tg^ mice failed to take up FM1-43 dye and to localize TMC1 at the stereocilia tips, indicating a mechanoelectrical transduction (MET) defect. Our data suggest a critical role of the cellular membrane to maintain functional hair cells and proper hearing and demonstrates that precautions must be considered when using these transgenic mice to study the auditory system, especially hair cell MET.

## MATERIALS AND METHODS

### Mice strains

6-8-week-old wild type C57BL/6J and homozygotes mT/mG mice (strains 000664 and 007676, respectively) were purchased from The Jackson laboratory and set to breed in our animal facility to obtain heterozygotes mT mice (mT/mG^Tg/+^). When the heterozygous progeny reached 6-8 weeks of age, female and male mT/mG^Tg/+^ mice were set to breed to obtain littermates of the three different phenotypes; mT/mG^Tg/Tg^, mT/mG^Tg/+^ and mT/mG^+/+^ for the mtdTomato. Genomic DNA extraction from tails snips and genotyping PCR reactions were performed using MyTaq Extract-PCR kit (Bioline, Taunton, MA). Mice were genotyped using a fragment PCR method with the primers shown in the table below and indicated on the Jackson Laboratory website. PCR products were run on a 2% agarose gel, and the Quick load 100pb DNA ladder (New England Biolabs Inc., Ipswich, MA) was used for fragment size visualization. PCR of the genomic wild type or mT/mG^+/+^ DNA gave a fragment of 212 bp, while genomic DNA from mT/mG^Tg/Tg^ gave a PCR fragment of 128 bp and the mT/mG^Tg/+^ mice gave both PCR fragments.

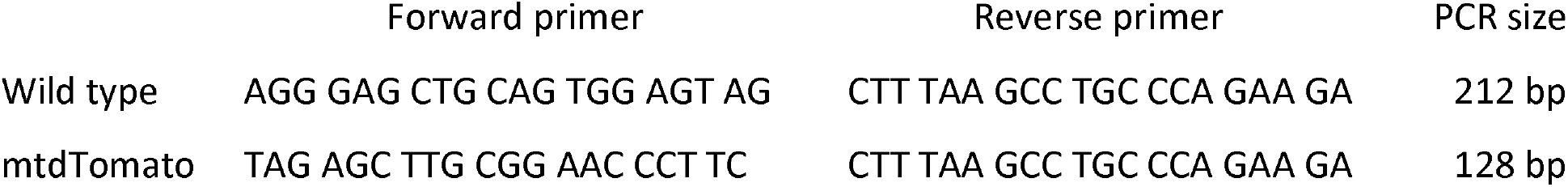

The mice were maintained at the AHCS animal facility under a 12:12 dark: light cycle, a temperature of 70-74°F and 35-60% humidity, and accessible NIH-07 rodent chow and water. The experiments performed with P6 mice included females and males since mice can’t be sexed at this age. The ABRs and DPOAEs recordings were performed in 3 males and 3 females for each genotype and no sex-related differences were observed. The animal care and experimental procedures were performed in accordance with the Guide for the Care and Use of Laboratory Animals and were approved by the Animal Care and Use Committee of the National Institute of Neurological Disorders and Stroke (animal protocol number 1336).

### Structural model of mtdTomato

A structural model of the mtdTomato fluorescent reporter was generated using I-TASSER (Yang et al., 2015). We used the protein sequence indicated below that includes the N-terminus of the MARCKS protein with two mutated cysteines (highlighted) that will result in the palmitoylation and myristoylation of the protein and its targeting to the plasma membrane. The x-ray structure of the fluorescent proteins mCerulean3 and cpVenus within the biosensor Twitch-2B (PDB ID: 6GEL) was used as a template (Trigo-Mourino et al., 2019). Template and query present a sequence coverage of the 83%, sequence identity of 25%, and 47.6% of sequence similarity. The palmitic acid and the myristoyl group structures were obtained from the PDB IDs 4OUG and 4ZV5. The POPC lipid bilayer was generated with the membrane builder tool of CHARMM-GUI with the default settings (Wu et al., 2014). The mtdTomato model was manually inserted in the membrane directed by the position of the lipidic motifs, and the final figure was generated with USCF Chimera (Pettersen et al., 2004). mtdTomato protein sequence:

> <u>MGCCFSKT</u> VSKGEEVIK EFMRFKVRME GSMNGHEFEI EGEGEGRPYE GTQTAKLKVT KGGPLPFAWD ILSPQFMYGS KAYVKHPADI PDYKKLSFPE GFKWERVMNF EDGGLVTVTQ DSSLQDGTLI YKVKMRGTNF PPDGPVMQKK TMGWEASTER LYPRDGVLKG EIHQALKLKD GGHYLVEFKT IYMAKKPVQL PGYYYVDTKL DITSHNEDYT IVEQYERSEG RHHLFLGHGT GSTGSGSSGT ASSEDNNMAV IKEFMRFKVR MEGSMNGHEF EIEGEGEGRP YEGTQTAKLK VTKGGPLPFA WDILSPQFMY GSKAYVKHPA DIPDYKKLSF PEGFKWERVM NFEDGGLVTV TQDSSLQDGT LIYKVKMRGT NFPPDGPVMQ KKTMGWEAST ERLYPRDGVL KGEIHQALKL KDGGHYLVEF KTIYMAKKPV QLPGYYYVDT KLDITSHNED YTIVEQYERS EGRHHLFLYG MDELYK

### DPOAEs and ABR recordings

Distortion product otoacoustic emissions (DPOAEs) and auditory brainstem responses (ABRs) were measured in littermate mice. All mice were anesthetized by intraperitoneal injection of 100 mg/kg ketamine (VetOne, MWI, Boise, ID, USA) and 0.4 mg/kg of DexDomitor (Dechra, Overlook Park, KS, USA) and placed inside a sound-attenuating booth (Acoustic Systems, ETS-Lindgren, Austin, TX, USA) on a heating pad connected to a temperature controller and rectal probe (ATC-2000, World Precision Instruments, Sarasota, FL, USA) to maintain body temperature near 37°C. Additional anesthesia was administered at one half the initial dose as necessary during testing.

Auditory tests were conducted using Tucker-Davis Technologies (TDT; Alachua, FL, USA) hardware (RZ6 Multi I/O processor, MF-1 speakers) and software (BioSigRZ, v.5.7.). DPOAEs were measured from both ears of each mouse using an ER-10B+ microphone (Etymotic, Elk Grove Village, IL, USA) connected to two MF-1 transducers (TDT). Two stimulus tones were presented at 14 frequency pairs spanning f2⍰=⍰4 to 40 kHz (L1⍰=⍰65 dB SPL and L2⍰=⍰55 dB SPL; f2/f1⍰=⍰1.2) and the amplitude of the DPOAE at 2f1 - f2 was recorded. Mean noise levels were calculated based on levels sampled at six surrounding frequencies. Mean emission and biological noise floor levels were calculated for each treatment group and plotted relative to each other.

ABR testing was conducted immediately after the DPOAE test was completed. Responses were recorded from subdermal needle electrodes (Rhythmlink, Columbia, SC, USA) placed at the vertex and under each pinna, with the non-test ear serving as the ground. Tone-burst stimuli (Blackman window, 3 ms duration, alternating polarity) were presented at a rate of 29.9/s via a closed-field TDT MF-1 speaker at 8, 16, 32, and 40 kHz. Responses were amplified (20x), filtered (0.3–3 kHz), and digitized (25 kHz). Threshold was determined by a visual inspection of stacked waveforms (512-1024 artifact-free responses per waveform). Stimuli at each frequency were presented initially at 80 dB SPL and decreased in 10-20 dB steps until the ABR waveform disappeared. The stimulus was then increased and decreased in 5 dB steps until the lowest sound intensity that resulted in an identifiable, repeatable waveform was determined. At least two waveforms were obtained for stimulus levels at and near the ABR threshold to ensure repeatability of the response. If no response was present at 80 dB SPL, stimulus level was increased to the maximum of 90 dB SPL and testing proceeded as described above. When no repeatable waves were visible at the highest stimulus level (90 dB SPL), the threshold was designated as 100 dB SPL for subsequent analyses. All threshold determinations were verified by an additional investigator.

When testing was complete, a subcutaneous injection of Antisedan (Zoetis) was injected to reverse the effects of anesthesia. Antisedan was dosed according to the NIDCD Veterinary and Husbandry Care Program guidelines based on body weight. Mice were placed in a cage devoid of bedding on a warming pad to fully recover before being returned to their original cage and to the animal facility.

Two orders of mT/mG mice were placed and received from The Jackson laboratory twice, in April 2018 and in September 2019. Homozygous mice from both orders presented no response at all the frequency tested indicative of profound deafness, including mT/mG shipped directly from The Jackson laboratory. Data from six mice (3 females and 3 males) received in September 2019 were included in this manuscript, and only data form one ear is shown for each mouse. Hearing was symmetrical in the mT/mG^+/+^ and mT/mG^Tg/Tg^ mice as both ears presented similar auditory phenotypes. However, mT/mG^Tg/+^ mice presented an asymmetrical hearing, especially at 16kHz, but the affected ear was not always the same one.

### Western Blots

Cochlear, vestibular, and cerebellar tissue were extracted from 3 months old littermates mT/mG^+/+^, mT/mG^Tg/+^ and mT/mG^Tg/Tg^ mice. As controls, liver and kidney tissue were extracted from the wild type mice. The tissue was washed with cold PBS buffer several times to remove blood and debris. Protein was extracted using cold T-PER buffer (ThermoFisher) supplemented with proteases inhibitors Complete Mini EDTA-free (ROCHE) and a rotor stator tissue homogenizer (OMNI). The tissue was kept on ice during the homogenization process. Homogenized samples were centrifuged for 30 min at 16000 rpm at 4 °C to remove the non-homogenized tissue, and supernatants were aliquoted and stored at −70 °C.

The total protein concentration in each sample was estimated by using the Pierce BCA protein assay kit (ThermoFisher). A standard curve for bovine serum albumin (BSA) was used to calculate the protein concentration of 1/10, 1/50, 1/100, and 1/1000 serial dilutions of the homogenized tissue samples in PBS. Equivalent amounts of total protein (50 ng/well) for each sample were loaded into a NuPAGE 4-12% Bis-Tris protein gel. Laemmli loading buffer supplemented with 100 mM DTT and 5% β-mercaptoethanol was added to the samples to a final concentration to facilitate the loading in the protein gel and to monitor the protein migration during the electrophoresis. Protein samples with loading buffer were heated at 70 °C for 10 min before loading into the gel. PageRuler Plus prestained protein ladder (ThermoFisher) was used to monitor the protein transfer on to the western blot membrane and to estimate the size of the proteins. NuPAGE MOPS SDS Running Buffer (ThermoFisher) was used as running buffer, and proteins were transferred to a nitrocellulose membrane by using the iBlot2 gel transfer device (ThermoFisher).

Membranes were blocked with 4% milk in Tris-Buffered Saline containing 0.1 % of tween 20 (TBST) buffer for 1h. Anti-prestin antibody (sc-293212, Santa Cruz) was added at 1/200 dilution in 4% milk in TBST buffer and incubated ON at 4 °C. Anti-TMC1 antibody (PB277) was used at 1/2000 in 4% milk in TBST buffer and incubated ON at 4 °C. Anti-RFP (660-401-379, Rockland) was added at 1/500 in 4% milk in TBST buffer and incubated ON at 4 °C. To detect the primary antibodies, a secondary anti-rabbit or anti-mouse antibody labeled with Horse Radish Peroxidase (HRP) (NA934VS, Millipore Sigma) was added at a 1/3000 dilution in 4% milk in TBST for 30 min at RT. As a protein loading control, the monoclonal anti-beta actin antibody labeled with HRP (ab20272, Abcam) was added in each western blot at 1/5000 dilution in 4% milk in TBST buffer and incubated for 30 min at RT. The blots were visualized and imaged in the ChemiDoc imaging system (BIO-RAD), and the Image Lab software (BIORAD) was used to quantify the amount of protein.

### Immunohistochemistry

Excision of the temporal bones and cochleae dissection, including removal of the semicircular canals and vestibular organs, were performed in Leibovitz’s L15 media (ThermoFisher) with surgical forceps under a stereomicroscope equipped with a WF10X eyepiece and an ACE light source. Cochleae were placed on a corning PYREX 9 depression plate well and washed with Hank’s balanced salt solution (HBSS, ThermoFisher). Tissue was fixed in 4 % paraformaldehyde (PFA, Electron Microscopy Science, Hatfield, PA) in HBSS buffer for 30 min. After incubation, tissue was washed twice with HBSS to remove PFA, and the spiral ligament and the tectorial membrane were both removed to obtain a fixed organ of Corti in HBSS buffer. Samples were then permeabilized in 0.5% Triton X-100 in PBS (Phosphate-Buffered Saline, ThermoFisher) for 30 min. Tissue was blocked with 4% BSA and 10% goat serum in PSB (blocking buffer) form 1h at RT and gentle shaking. Anti-prestin antibody (sc-293212, Santa Cruz), antimyosin VIIa antibody (25-6790, Proteus), or anti TMC1 antibody (PB277) were added in blocking buffer at a dilution of 1/200, 1/500 or 1/1000, respectively, and incubated ON at 4°C with gentle shaking. The next morning, tissue was washed 2-3 times with PBS buffer. Secondary antibody at a 1/1000 dilution (Alexa Fluor 594 goat anti-rabbit antibody or Alexa Fluor 488 goat anti-mouse antibody, Thermo Fisher) in blocking buffer also containing Alexa Fluor-405 phalloidin (ThermoFisher Scientific) at a 1:200 dilution for 30 min was added and incubated for 1h at RT. Tissue was washed with PBS at least 3 times and was finally mounted with ProLong Diamond antifade mounting media (ThermoFisher Scientific) on a superfrost plus microscope slide (Fisherbrand, Pittsburgh, PA) and covered with a #1.5 glass coverslips of 0.17 ± 0.02 mm thickness (Warner Instruments, Hamden, CT) for confocal imaging.

### FM1-43 dye uptake experiments

Excision of the temporal bones from littermates mT/mG^+/+^, mT/mG^Tg/+^ and mT/mG^Tg/Tg^ P6 mice and further cochleae dissection, including removal of the semicircular canals and vestibular organs, were performed in Leibovitz’s L15 media with surgical forceps under an amplification stereomicroscope. Two incisions were performed on the dissected cochleae free of surrounding tissue, one on the round window and other at the apical cochlear region. Cochleae were placed on a corning PYREX 9 depression plate well in HBSS buffer and incubated for 2 min at room temperature with FM1-43 dye at 5 μM in HBSS with gentle shaking. After incubation, tissue was washed 5 times with HBSS, and fixed in 4 % paraformaldehyde (PFA, Electron Microscopy Science, Hatfield, PA) in HBSS for 30 min. Fixed tissue was washed with HBSS to remove PFA. The spiral ligament and the tectorial membrane were both removed to obtain fixed organ of Corti in HBSS buffer. Tissues were washed 2-3 times with PBS buffer to and were finally mounted with ProLong Diamond antifade mounting media (ThermoFisher Scientific) on a superfrost plus microscope slide (Fisherbrand, Pittsburgh, PA) and covered with a #1.5 glass coverslips of 0.17 ± 0.02 mm thickness (Warner Instruments, Hamden, CT) for confocal imaging.

### Image acquisition

Imaging was performed in the Microscopy and Imaging Core (NICHD) with a confocal laser scanning microscope Zeiss LSM 880 (Carl Zeiss AG, Oberkochen, Germany) equipped with a 32 channel Airyscan detector (Korobchevskaya et al., 2017). To image the hair cells, we collected a z-stack of images from the stereocilia to the apical half of the hair cell body. We used oil immersion alpha Plan-Apochromat 63X/1.4 Oil Corr M27 objective (Carl Zeiss) and Immersol 518F immersion media (ne=1.518 (30°), Carl Zeiss). Identical image acquisition settings, no averaging, and optimal parameters for x, y, and z resolution were used in all samples from each independent experiment. Image acquisition and Airyscan image processing were made with Zen Black 2.3 SP1 software (Carl Zeiss) using the Airyscan 3D reconstruction algorithm with the automatic default Wiener filter settings. The images of the whole organ of Corti samples were captured with a 20X objective, and a z-stack of 3 confocal planes was captured for each organ. The tiles were stitched with a 10% overlay, and a maximum intensity projection image was generated using the Zen software. Confocal imaging on FM1-43 uptake experiments and whole organ of Corti samples was performed in the Microscopy and Imaging Core (NICHD) with an inverted laser scanning microscope Zeiss LSM 780 (Carl Zeiss) equipped with a motorized stage, definite focus and a high sensitivity GaAsp multi-channel spectral detector. To image these samples, we used a 63X/1.4 objective Plan-Apochromat (Carl Zeiss) and the Zen software (Carl Zeiss). Identical image acquisition settings, no averaging, and optimal parameters for x, y, and z resolution were used in all samples from each independent experiment. Representative confocal images for each condition in a representative experiment are shown with the same display range.

### Data processing and statistical analysis

Microscopy data processing, analysis, and quantification were done in ImageJ (Schneider et al., 2012). To measure the fluorescence intensity at the cochlear regions, we generated 10 regions of interest (ROI) for each region of the cochlea (basal, middle, and apical) along the organ of Corti using the oval tool and measured the integrated mean fluorescence intensity of each ROI. Three independent organs of Corti preparations from littermates mT/mG^+/+^, mT/mG^Tg/+^ and mT/mG^Tg/Tg^ mice were used. ROI were generated at each hair cell at the cell body or stereocilia plane for the quantification of prestin or TMC1 fluorescent intensity. An equivalent ROI was generated at a region outside the hair cells, considered as background and subtracted from the hair cell fluorescent intensity. Background subtraction was performed using a rolling ball of 200 pixels for the figures shown in the manuscript.

Data were initially processed in Microsoft Excel, and GraphPad Prism V.7 software (GraphPad Software, La Jolla, CA) was used to generate the graphs and perform the statistical analysis. Ordinary one-way ANOVA analysis with multiple comparisons of the mean was performed for all the quantifications reported. We used the style GP to represent the p-value style GP, which was indicated in the figures as p<0.1234 as no significative (n.s.), p<0.0332 as *, p<0.0021 as **, p<0.0002 as *** and p<0.0001 as ****. Analysis of the fluoresce intensity were corrected for multiple comparisons using Tukey test, while statistical analysis of the western blot triplicates was not corrected for multiple comparisons and performed using the Fisher’s LSD test.

## RESULTS

### The mT/mG mouse as a tool to visualize the auditory hair cell membrane

mT/mG mice carry a transgene encoding two fluorescent tomato proteins in tandem with the first 8 amino acids of the Myristoylated Alanine-Rich C-Kinase Substrate (MARCKS) with the third and fourth amino acids mutated to cysteines (MGCCFSKT) at its N-terminus (Muzumdar et al., 2007). The endogenous MARCKS is myristoylated on the N-terminal glycine (Stumpo et al., 1989), whereas the introduced two cysteines act as palmitoylation sites (Wiederkehr et al., 1997) that direct mtdTomato to the cell membrane. Representation of an approximated model of a mtdTomato protein in a lipid membrane suggested that each reporter would occupy a relatively small region of the intracellular side of the membrane (Figure 1B).

To visualize the membrane of the auditory hair cells in mT/mG^Tg/+^ mice at high resolution, we used a Zeiss LSM 880 confocal microscope equipped with an Airyscan detector (see Methods), and imaged organ of Corti explants of mT/mG^Tg/+^ P6 mice. The mtdTomato reporter labeled the stereocilia membrane of both the inner hair cells (IHC) and the outer hair cells (OHC), which the latter presenting a more intense labeling (Figure 1C). The kinocilial vestigial area, known to present intense vesicular trafficking (Kachar et al., 1997), was highlighted by the expression of the mtdTomato reporter (Figure 1C, yellow arrows in the top right panel). We also observed several membranous organelles below the cuticular plate that were absent when approaching the nuclear region (Figure 1C, yellow arrows in the bottom left panel). The apical and basal plasma membrane of both types of hair cells was also visible in mT/mG mice (Figure 1C).

### mT/mG^Tg/Tg^ mice are profoundly deaf

mT/mG^Tg/Tg^, mT/mG^Tg/+^ and mT/mG^+/+^ littermates had no observable phenotype, including no gross vestibular anomalies such as circling behavior or head bobbing, suggesting minimal toxicity by the expression of the mtdTomato reporter *in vivo*. However, upon weaning, it was apparent that some offspring lacked the Preyer reflex, the ear flick response to sound (Jero et al., 2001), suggesting hearing impairment. To test for hearing deficits, we characterized the auditory phenotype of mT/mG^Tg/Tg^ and mT/mG^Tg/+^ mice by measuring auditory brainstem responses (ABRs) and distortion product otoacoustic emissions (DPOAEs) (Avan et al., 2013; Brownell, 1990; Skoe et al., 2010).

At 6-8 weeks of age, mT/mG^Tg/+^ mice presented elevated ABR thresholds at 32 and 40 kHz compared to mT/mG^+/+^ littermates, indicating a high-frequency hearing loss (Figure 2A). More remarkably, mT/mG^Tg/Tg^ littermate mice did not respond to even the highest levels of sound (90 dB SPL), indicating profound deafness at all frequencies tested (Figure 2A). We also measured DPOAEs evoked by low-level tones, which are an indirect measure of outer hair cell (OHC) mechanical function (Shaffer et al., 2003). mT/mG^Tg/+^ mice presented elevated DPOAEs amplitudes at the highest frequency tested compared to their mT/mG^+/+^ littermates (Figure 2B), indicating compromised OHC function at the basal region of the cochlea, where the high frequencies are transduced. DPOAEs above 20kHz (noise level) were absent in all the mT/mG^Tg/Tg^ mice tested, in agreement with the ABR test results and indicating severe loss of OHC function (Figure 2B). Overall, these data indicate that the expression of the mtdTomato reporter impairs the function of auditory OHC at the higher frequencies in mT/mG^Tg/+^ mice and all frequencies in mT/mG^Tg/Tg^ mice.

**Figure 2:**
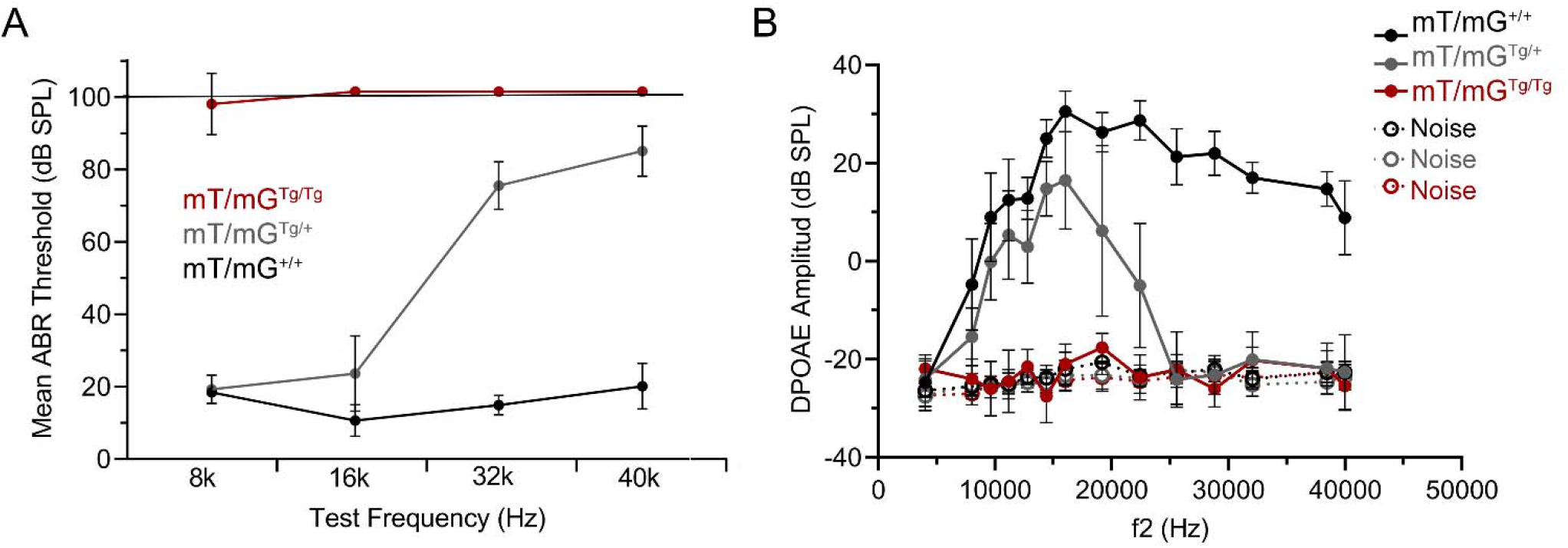
ABR and DPOAEs thresholds are elevated in the mT/mG reporter mice. **A)** Average auditory brainstem response (ABR) thresholds in littermate mT/mG^+/+^(red), mT/mG^Tg/+^ (grey) and mT/mG^Tg/Tg^ (black) mice. ABRs were measured in both ears of 6 (3 females and 3 males) from each of the three genotypes at frequencies of 8, 16, 32 and 40 kHz in 40-60 days old mice. When no response was detected at a maximum stimulus level of 90 dB SPL, the threshold was assigned as 100 dB SPL (dashed black line). Error bars indicate SD. **B)** DPOAE amplitudes measured in the same mT/mG^+/+^(red), mT/mG^Tg/+^ (grey), and mT/mG^Tg/Tg^ (black) mice shown in A. Mean noise floors were calculated for all mice using six samples, three on either side of the DPOAE frequency, and are indicated by the dashed lines in the same color that its corresponding phenotype (noise). Error bars indicate SD.

### Expression of the mtdTomato reporter in the organ of Corti of young adult mice

A significant loss of OHCs is commonly associated with a reduction in the DPOAE amplitudes (Fettiplace et al., 2019; Ryan et al., 1980; Salvi et al., 2016). To assess if the hearing phenotype of the mT/mG^Tg/+^ and mT/mG^Tg/Tg^ mice was due to the loss of OHCs, we examined the presence of OHC in adult mice. We found that the three rows of OHC were present at the middle-apical area of the organ of Corti from 47 days old mT/mG^Tg/Tg^, mT/mG^Tg/+^ and mT/mG^+/+^ littermates and the hair cell bundle and stereocilia were comparable in all three genotypes (Figure 3A and B). We concluded that the presence and overall morphology of hair cells and OHC are preserved in adult mT/mG mice.

**Figure 3:**
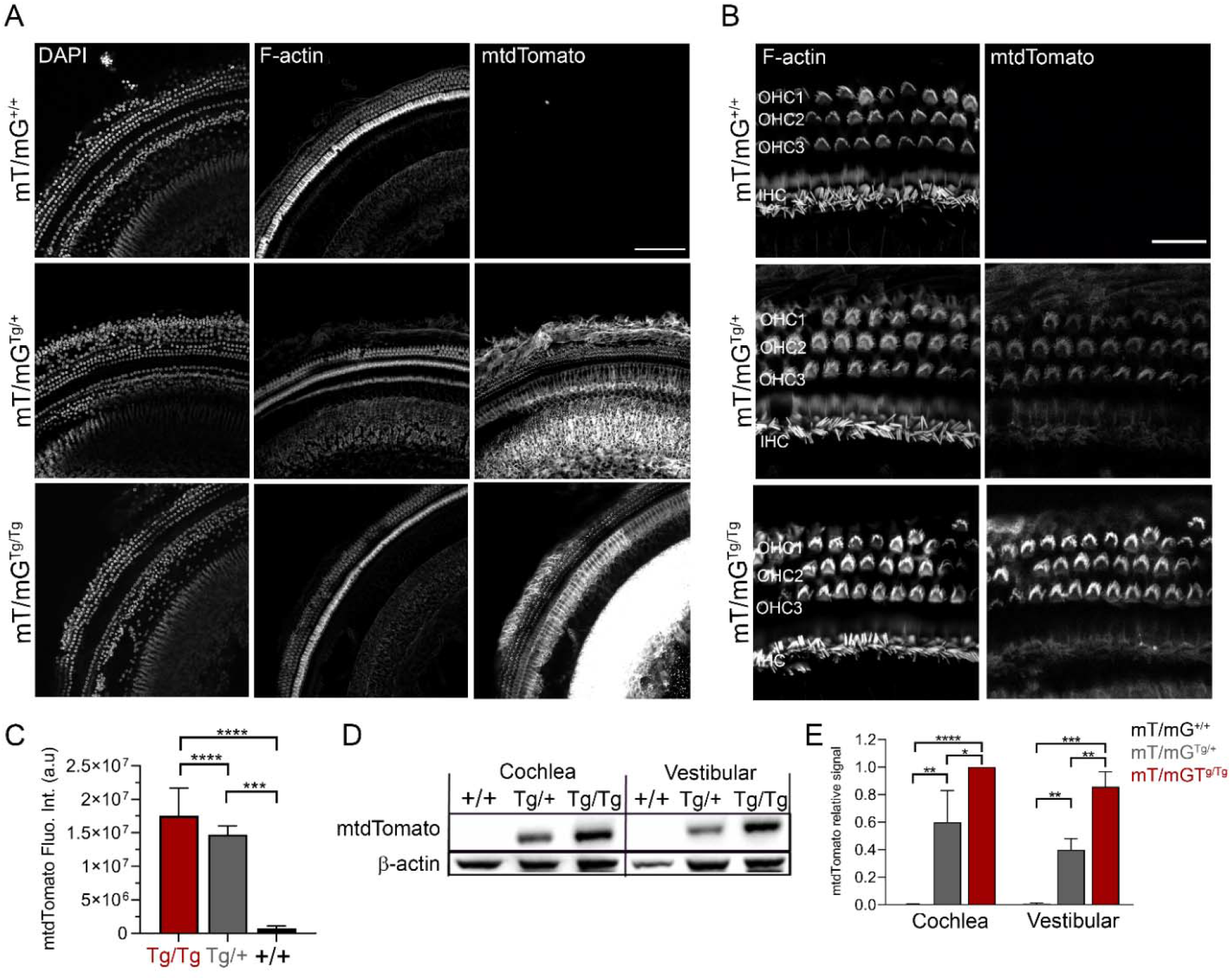
Expression of the mtdTomato transgene in the adult cochleae. **A)** Middle region of the organ of Corti of mT/mG^+/+^, mT/mG^Tg/+^ and mT/mG^Tg/Tg^ P47 adult mice. The nucleus of the hair cells was labeled with DAPI and phalloidin was used to label the F-actin filaments and visualize the stereocilia of the hair cells. mtdTomato fluorescent signal at the apical cochlear region is also shown. Scale bar represents 100 μm. **B)** Close up view of the hair cells from the middle-apical region of the cochlea. The three rows of OHC and the one row of IHC are labeled. Scale bar represents 20 μm. **C)** Quantification of the fluoresce intensity of the mtdTomato reporter at the sensory hair cells region of the middle-apical area of the cochlea. **D)** Western blot for the detection of the expression levels of the mtdTomato protein at the cochlea and vestibular organs mT/mG^+/+^, mT/mG^Tg/+^ and mT/mG^Tg/Tg^ mice. mtdTomato was detected using an anti-RFP antibody. An antibody against β-actin was used to control the amount of protein loaded. **E)** Quantification of the mtdTomato signal from western blots relative to the tissue with the highest protein level and normalized against the total loaded protein (β-actin). Values are mean ± SD from 3 experiments. One-way ANOVA analysis was performed in C and E (*p<0.0332, **p<0.0021, ***p<0.0002, and ****p<0.0001).

Quantification of the fluorescent intensity of the mtdTomato at the hair cell area, including OHC and IHC, revealed that the fluorescent signal was similar to background levels in mT/mG^+/+^ mice and stronger in mT/mG^Tg/Tg^ mice than in mT/mG^Tg/+^ (Figure 3C). The expression levels of mtdTomato in the 3 different genotypes was further confirmed by western blot analysis of cochlear and vestibular tissue samples (Figure 3D and 3E). Our confocal and western blot data indicate that the reporter is expressed at higher levels in the mT/mG^Tg/Tg^ than in mT/mG^Tg/+^ adult mice.

### Expression of the mtdTomato reporter in the organ of Corti of neonate mice

The study of the cochlea in adult animals becomes difficult due to calcification of the cochlea, making the dissection of the basal and middle regions of the organ of Corti challenging. Therefore, to evaluate the expression of the mtdTomato along the cochlea, including its basal region, we performed a similar analysis in neonatal mice.

We first inspected the presence of auditory IHC and OHC along the cochlea. As shown in Figure 4A, OHC and IHC located at the apical, middle, and basal areas of the cochlea were present at this age. We also examined the mtdTomato transgene expression at the hair cells in mT/mG^Tg/Tg^, mT/mG^Tg/+^ and mT/mG^+/+^ P6 littermate mice. The mtdTomato fluorescence was absent in mT/mG^+/+^ mice (Figure 4A and B), and we observed a higher fluorescence intensity of the reporter in the mT/mG^Tg/Tg^ mice compared to the mT/mG^Tg/+^ (Figure 4A-C), similar to that observed in adult mice. Interestingly, we found an increase in the fluorescence intensity at the basal area of the cochlea compared to the apical area in mT/mG^Tg/+^ and mT/mG^+/+^ mice (Figure 4B). Importantly, we did not perceive any morphological differences between the hair cells of mT/mG^Tg/Ts^, mT/mG^Tg/+^ and mT/mG^+/+^ mice at the stereocilia level (Figure 4D). The organization of the stereocilia, overall bundle architecture, and bundle orientation of the OHC and IHC were comparable in all genotypes (Figure 4D).

**Figure 4:**
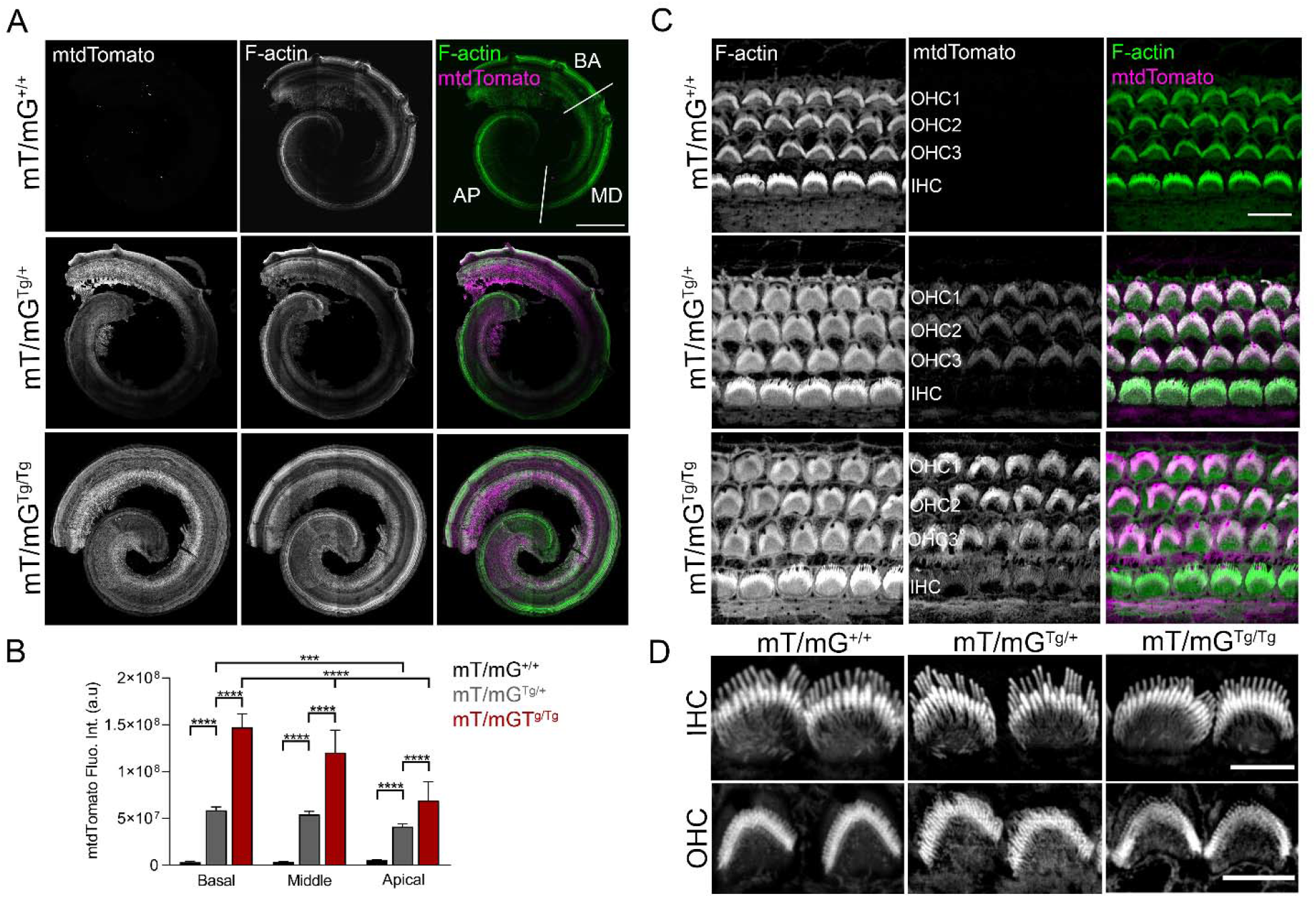
Expression of the mtdTomato transgene in the neonate cochleae. **A)** Confocal images of the whole organ of Corti from P6 mT/mG^+/+^, mT/mG^Tg/+^ and mT/mG^Tg/Tg^ mice. Images from the F-actin labeling and the mtdTomato reporter are shown in grey independently and merged in green (F-actin) and magenta (mtdTomato). Scale bar represents 300 μm. **B)** Quantification of the fluoresce intensity of the mtdTomato at the sensory hair cells region of the basal (BA), middle (MD), and apical (AP) areas of the cochlea of the mT/mG^+/+^(black), mT/mG^Tg/+^ (grey) and mT/mG^Tg/Tg^ (red) mice. Mean fluorescence intensity ± SD is represented. One-way ANOVA analysis was performed (***p<0.0002, and ****p<0.0001). **C)** Confocal images of hair cells from the middle area of the cochlea of a P6 mouse represented as in A. Scale bar represents 10 mm. The three rows of OHC and the one row of IHC are labeled. **D)** IHC (top panel) or OHC (lower panel) from mT/mG^+/+^, mT/mG^Tg/+^ and mT/mG^Tg/Tg^ mice stained for phalloidin. Scale bar represents 5 μm.

### Prestin expression and localization is preserved in the mT/mG mice

We wondered whether the expression of the mtdTomato reporter could affect one of the principal characteristics of OHC, their somatic motility. OHC change the length and axial stiffness of their cell bodies in response to changes in their membrane potential (Brownell, 1990; Brownell et al., 1985; He et al., 2010). This electromotility, essential for cochlear amplification and DPOAEs, is driven by the molecular motor prestin, which is expressed at the lateral plasma membrane of mature OHC and absent from IHC (Abe et al., 2007; Adler et al., 2003; He et al., 2010; Takahashi et al., 2018).

To evaluate if the expression of the mtdTomato transgene interferes with the expression on prestin in OHC, we analyzed by western blot the expression of prestin in the cochlea and vestibular system of littermates mT/mG^Tg/Tg^, mT/mG^Tg/+^ and mT/mG^+/+^adult mice (Figure 5A) and found that all three genotypes expressed similar levels of prestin (Figure 5B). However, the transgene could disturb the localization of the motor protein prestin without affecting its overall expression levels. To test this, we localized prestin by immunohistochemistry in the organ of Corti from P10 mice, when prestin expression can be detected (Belyantseva et al., 2000). We detected the expression of prestin at the lateral plasma membrane of auditory OHC in the organ of Corti of mT/mG^Tg/Tg^, mT/mG^Tg/+^ and mT/mG^+/+^ mice (Figure 5C). Quantification of the immunofluoresce intensity of prestin in the mT/mG^Tg/Tg^, mT/mG^Tg/+^ and mT/mG^+/+^ mice revealed similar levels of prestin in the three genotypes (Figure 5D). We concluded that the expression levels and localization of prestin are not affected by mtdTomato in mT/mG mice.

**Figure 5:**
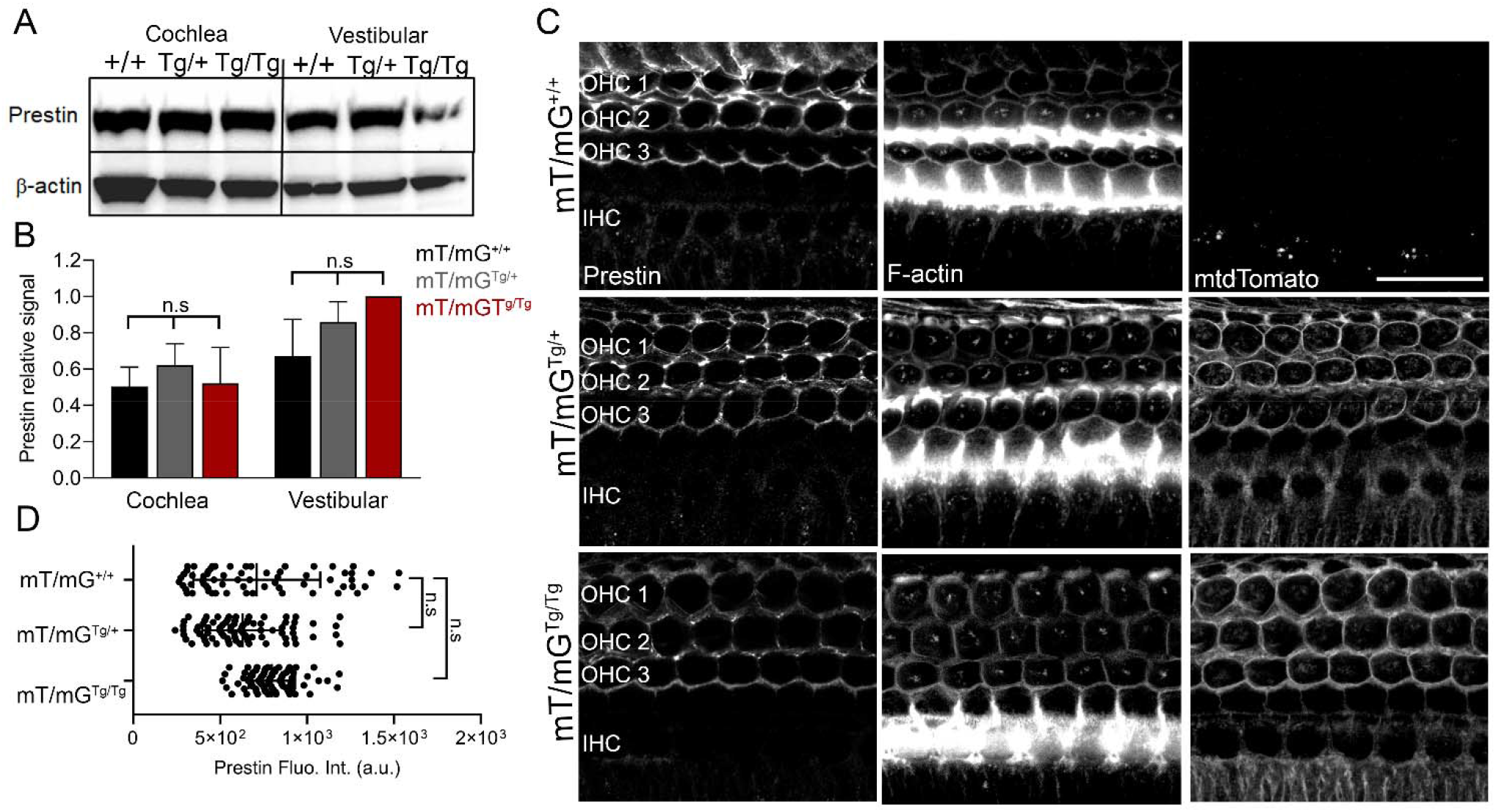
Expression and localization of prestin is preserved in mT/mG mice. **A)** Representative western blot of the prestin levels at the cochlea and vestibular organs of mT/mG^+/+^, mT/mG^Tg/+^ and mT/mG^Tg/Tg^ mice. An antibody against β-actin was used to control the amount of protein loaded. **B)** Quantification of the prestin signal relative to the tissue with the highest prestin level and normalized against the total loaded protein (β-actin). Values are mean ± SD from 3 blots. **C)** Representative confocal images of hair cells of P10 mT/mG^+/+^, mT/mG^Tg/+^ and mT/mG^Tg/Tg^ mice. Fluorescent intensity from prestin, F-actin labeling or the mtdTomato reporter are shown in grey independently. The three rows of OHC and the one row of IHC are labeled. Scale bar represents 20 μm. **D)** Quantification of the fluorescence intensity of prestin at the OHC body in the mT/mG^+/+^, mT/mG^Tg/+^ and mT/mG^Tg/Tg^ mice. Mean fluorescence intensity ± SD is represented for n = 65-80 hair cells. One-way ANOVA analysis was performed in B and D (n.s. p<0.1234).

### Mechanoelectrical transduction is altered in mT/mG^Tg/Tg^ mice

We next evaluated whether the expression of the mtdTomato reporter alters the MET channel activity in mT/mG^Tg/Tg^ and mT/mG^Tg/+^ mice by measuring the uptake of the styryl dye FM1-43. The FM1-43 dye is commonly used to assay the presence of MET channel activity at rest since functional MET channels are required for the uptake of this fluorescent dye into hair cells (Gale et al., 2001; Kawashima et al., 2015; Meyers et al., 2003). We exposed organ of Corti explants from P6 mice to a brief incubation with FM1-43 and observed a robust uptake of the fluorescent dye into hair cells from mT/mG^+/+^ and mT/mG^Tg/+^ mice. However, hair cells from mT/mG^Tg/Tg^ littermate mice failed to uptake FM1-43 (Figure 6), suggesting that the MET channel activity is compromised in neonatal mT/mG^Tg/Tg^ mice.

**Figure 6:**
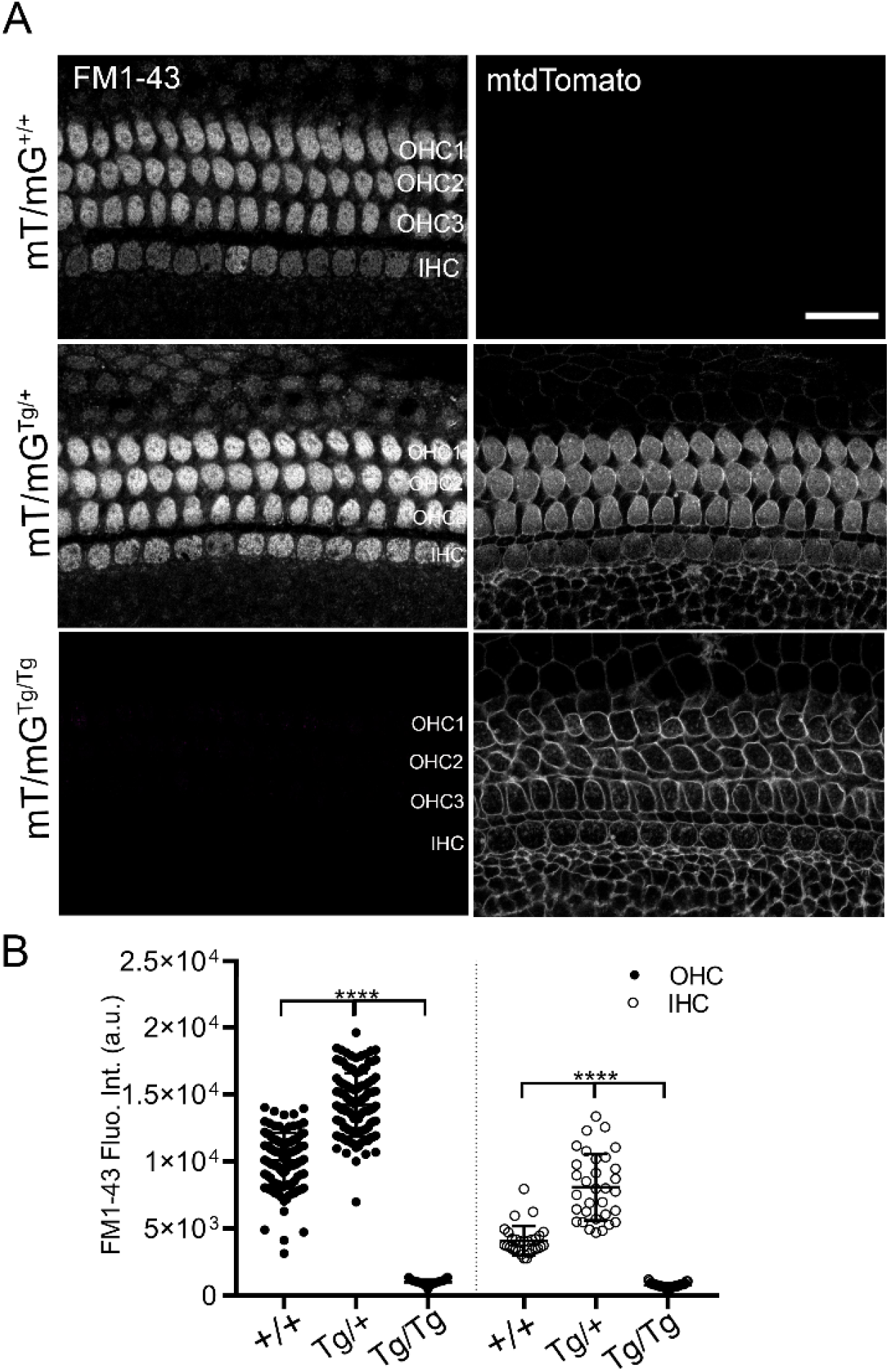
Hair cells form mT/mG^Tg/Tg^ mice fail to uptake the FM1-43 dye. **A)** Representative confocal images of the FM1-43 uptake in hair cells of mT/mG^+/+^, mT/mG^Tg/+^ and mT/mG^Tg/Tg^ P6 mice. Fluorescent intensity from the mtdTomato and FM1-43 dye channels are shown in grey independently. The three rows of OHC and the one row of IHC are labeled. Scale bar represents 20 μm. **B)** Quantification of the FM1-43 fluorescence intensity at the hair cell body of OHC (⍰) and IHC (⍰) of mT/mG^+/+^, mT/mG^Tg/+^ and mT/mG^Tg/Tg^ mice. Mean fluorescence intensity ± SD is represented. The number of hair cells analyzed in each condition (n) is 110, 116, 110, 32, 29 and 33One-way ANOVA analysis was performed in B and D (****p<0.0001).

Increasing evidence supports the involvement of the TransMembrane Channel-like protein 1 (TMC1) in forming the pore of the MET channel complex at the tips of hair cell stereocilia (Beurg et al., 2015; Beurg et al., 2019; Corns et al., 2017; Jia et al., 2020; Kawashima et al., 2015; Kawashima et al., 2011; Kurima et al., 2015; Pan et al., 2013; Pan et al., 2018). To investigate whether the mtdTomato reporter alters the expression of TMC1, we analyzed the expression of TMC1 in cochlear and vestibular tissue from mT/mG^Tg/Tg^, mT/mG^Tg/+^and mT/mG^+/+^mice by western blot. Similar TMC1 protein levels were detected in the cochlea and vestibular organs from mT/mG^Tg/Tg^, mT/mG^Tg/+^ and mT/mG^+/+^ mice (Figures 7A-B). However, immunolabeling of TMC1 in the organ of Corti showed that TMC1 was absent from the stereocilia of hair cells from mT/mG^Tg/Tg^ mice. To explore whether the localization of TMC1 is altered by expression of mtdTomato, we detected the expression of TMC1 in the organ of Corti by immunohistochemistry. Remarkably, immunolabeling of TMC1 in the organ of Corti showed that TMC1 was absent from the stereocilia of the hair cells from mT/mG^Tg/Tg^ mice, whereas it localized to the stereocilia in mT/mG^+/+^ and mT/mG^Tg/+^ mice (Figure 7C-D). Therefore, we concluded that mtdTomato, when expressed at high levels, alters the proper localization of one of the main components of the MET complex.

**Figure 7:**
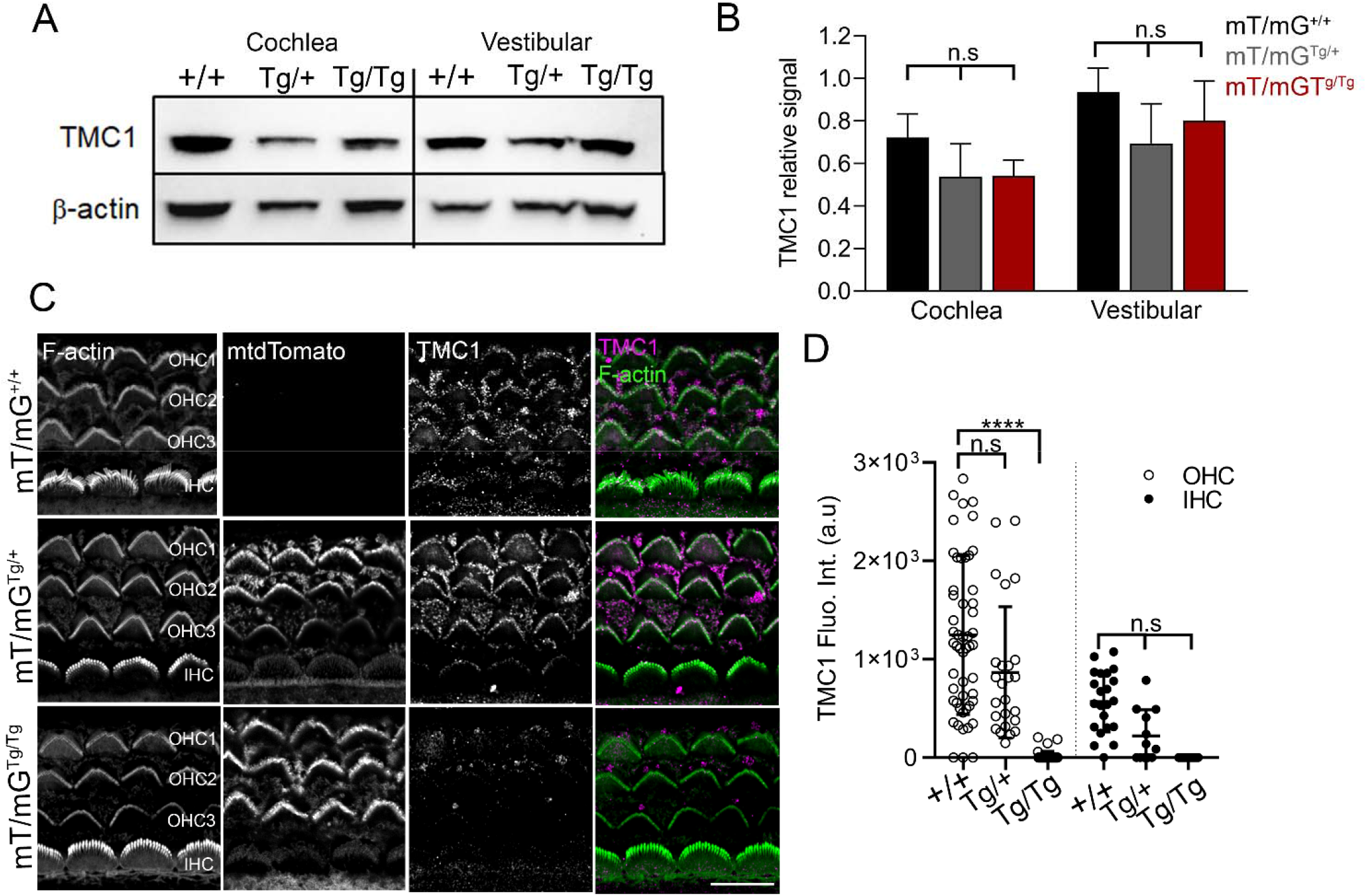
Hair cells form mT/mG^Tg/Tg^ mice fail to localize TMC1 at the stereocilia tips. **A)** Representative western blot of the TMC1 levels at the cochlea and vestibular organs of mT/mG^+/+^, mT/mG^Tg/+^ and mT/mG^Tg/Tg^ mice. An antibody against β-actin was used to control the amount of loaded protein. **B)** Quantification of the TMC1 signal relative to the tissue with the highest TMC1 level (vestibular mT/mG^+/+^) and normalized against the total loaded protein (β-actin). Values are mean ± SD for 3 blots. **C)** Immunostaining of basal hair cells of the organ of Corti of P6 mT/mG^+/+^, mT/mG^Tg/+^ and mT/mG^Tg/Tg^ mice with an antibody anti-TMC1 (PB277). The F-actin, TMC1 and tomato channels are shown independently in gray. The merged F-actin (green) and TMC1 (magenta) channels are also shown. Scale bar is 10 μm. **D)** Quantification of the TMC1 immunofluorescence intensity at the hair cell stereocilia of OHC (⍰) and IHC (⍰) of mT/mG^+/+^, mT/mG^Tg/+^ and mT/mG^Tg/Tg^ mice. Background-subtracted mean fluorescence intensity ± SD is represented. The number of hair cells analyzed in each condition and represented as a dot (n) is 53, 42, 25, 22, 16, 8. One-way ANOVA analysis was performed in B and D (n.s. p<0.1234 and ****p<0.0001).

## DISCUSSION

ROSA26 is assumed to be a safe locus for the generation of viable and fertile mouse reporter lines with unaltered phenotypes, which appears to justify the fact that these lines are typically encouraged to be maintained and used in a homozygous state (Li et al., 2018; Muzumdar et al., 2007; Soriano, 1999; Srinivas et al., 2001; Zambrowicz et al., 1997). However, we demonstrate that although mT/mG^Tg/Tg^, mT/mG^Tg/+^ and mT/mG^+/+^ littermates are viable, fertile, and indistinguishable in appearance, the expression of the mtdTomato reporter gene inserted in the ROSA26 locus severely affects the auditory function of adult mT/mG^Tg/Tg^ and mT/mG^Tg/+^ mice as assessed by ABR and DPOAE recordings. mT/mG^Tg/Tg^ mice presented profound hearing loss at all the frequencies tested, and mT/mG^Tg/+^ mice showed a severe high-frequency hearing loss at 32 and 40 kHz when compared to mT/mG^+/+^ littermates. DPOAEs were absent in the high frequencies for the mT/mG^Tg/+^ mice and absent at all frequencies in the mT/mG^Tg/Tg^ mice, consistent with the ABRs, revealing a high-frequency hearing loss and profound hearing loss at all frequencies in the mT/mG^Tg/+^ and mT/mG^Tg/Tg^ mice, respectively (Figure 2). Our data indicate that the expression of the reporter transgene in the auditory system affects the function of OHC and the ability of the transgenic mice to perceive sound. Consistent with this, the higher expression levels of mtdTomato in mT/mG^Tg/Tg^ adult and neonatal mice (Figure 3 and 4) correlated with a more severe auditory deficiency in these mice.

DPOAEs are produced when stimulus tones are delivered to the inner ear and interact with the basilar membrane to stimulate nonlinear elements in the cochlea that amplify the acoustic stimuli. OHC electromotility is the primary source of this ‘cochlear amplifier’. Prestin is responsible for the OHC electromotility and localizes to the plasma membrane of the lateral wall region of OHC (Cortese et al., 2017; Yamashita et al., 2015). However, we show here that the expression and localization of prestin are comparable in mT/mG^Tg/Tg^, mT/mG^Tg/+^ and mT/mG^+/+^ littermate mice (Figure 5). Furthermore, since OHCs prematurely deteriorate and die in prestin knock-out mice or transgenic mice expressing a non-functional prestin, the lack of hair cell loss in mT/mG mice supports a marginal role of prestin in the auditory phenotype of these mice (Cheatham et al., 2015; Liberman et al., 2002). Therefore, while we cannot rule out that the electromotile function of prestin is affected by the expression of the mtdTomato transgene, the preservation of OHCs and the expression and localization of prestin in mT/mG mice suggest that a different mechanism underlies the auditory defect of these mice.

MET, the process through which cells sense and respond to mechanical stimuli by converting them to electrochemical signals, is the signature function of hair cells. Hair cells that lack MET fail to uptake small styryl dyes, such as FM1-43(Gale et al., 2001; Kawashima et al., 2011; Park et al., 2013). Here we show that hair cells from mT/mG^Tg/Tg^ mice are unable to uptake FM1-43, indicating that these mice lack functional MET channels that are open at rest (Figure 6) (Gale et al., 2001). This observation suggests an important role of the cell membrane in regulating the MET channel, which has been previously suggested (Effertz et al., 2017; Gianoli et al., 2017; Hirono et al., 2004). TMC1 and TMC2 proteins are thought to contribute to forming the pore of the MET channel and are essential for MET (Jia et al., 2020; Kawashima et al., 2015; Kawashima et al., 2011; Pan et al., 2013; Pan et al., 2018). Consequently, hair cells from TMIE and mTOMT knock out mice that lack MET, fail to properly localize TMC1 and TMC2 at their stereocilia tips (Cunningham et al., 2017; Cunningham et al., 2020). Although we did not specifically analyze TMC2 because the expression of TMC2 is transient in early postnatal murine cochlear hair cells and TMC1 is responsible for auditory MET in mature hair cells, the absence of TMC1 and TMC2 at the stereocilia in the mT/mG^Tg/Tg^ mice explains the lack of MET in these mice (Figure 7). The stereocilia bundle of hair cells from TMC1 knock out and TMC1/TMC2 double knock out mice exhibits an underdeveloped phenotype consisting of a U-shaped bundle with multiple rows of stereocilia, and some OHC loss in the middle and basal cochlear regions has been reported in these knock out mice at P25 (Kawashima et al., 2011; Lefevre et al., 2008; Nakanishi et al., 2018). However, the overall hair cell bundle architecture and stereocilia organization of the mT/mG^Tg/Tg^ mice was comparable to the mT/mG^+/+^, and we did not perceive OHC loss at P47, suggesting a milder and slower deteriorating phenotype. The preserved expression levels of TMC1 in the mT/mG^Tg/Tg^ mice could explain this milder phenotype and indicates that the expression of the mtdTomato reporter interferes the folding, trafficking or the stability of TMC1 at the hair cell stereocilia.

The mT/mG^Tg/Tg^ mice do not present a vestibular phenotype, suggesting that the expression of the mtdTomato reporter only affects the function of the auditory hair cells. Interestingly, sensory transduction is maintained throughout the utricle in the absence of TMC1 (Asai et al., 2018; Corns et al., 2017), and TMC1 knock-out mice are deaf but present no vestibular phenotype, suggesting a minor role for this protein in vestibular hair function (Kawashima et al., 2015). Moreover, the function of other proteins known to be essential for hair cell MET (TMIE, LHFPL5/TMHS, MYO7a, CDH23, and PCDH15) may not be affected by the expression of the mtdTomato reporter since mutation of these genes lead to both auditory and vestibular deficits in humans and mouse models (Ahmed et al., 2001; Lefevre et al., 2008; Longo-Guess et al., 2005; Park et al., 2013).

Neonatal mT/mG^Tg/+^are an appropriate tool to study and visualize the neonate hair cell membrane, and mT/mG mice have been used in several studies in the auditory field. For instance, mT/mG^Tg/+^ crossed with Slc26a5-CreER^T2^ have been used to look at the lateral diffusion of membrane proteins in OHCs (Yamashita et al., 2015). Consistent with our data, these mice exhibited elevated ABR thresholds at 44kHz at P20. Interestingly, mT/mG^f/+^ mice expressing mGFP following Cre recombination also exhibited elevated ABR thresholds at mid-high frequencies (22–32 kHz) (Yamashita et al., 2015), suggesting that the expression of the mGFP transgene, which is a monomeric protein but has a 33-residues longer MARCK N-termini, could affect the auditory function more dramatically. Although generally thought to not interfere with the membrane properties and microdomains (Abe et al., 2013; Laguerre et al., 2018; Sezgin et al., 2017; Varnai et al., 2007), the expression of fluorescent reporters near the cell membrane could hinder the interaction of cytosolic or membrane proteins with the plasma membrane. For instance, mtdTomato could interfere with the interaction of TMIE with phosphatidylinositol lipids, which may be necessary for the proper localization of TMC1 at the stereocilia (Cunningham et al., 2020). Although the viability and lack of a visible phenotype suggest that the function of other mechanosensitive ion channels may be preserved in the mT/mG mice, it would be interesting to study if other physiological processes associated with mechanosensory transduction are also affected, or why the expression of this reporter specifically affects the localization of TMC1 and the activity of the auditory hair cell MET channel. Until then, we recommend using neonatal mT/mG^Tg/+^and avoiding the mT/mG^Tg/Tg^ mice to study auditory hair cell functions.

## ACKNOWLEDGMENTS

We thank Dr. Vincent Schram from the NICHD microscopy and imaging core for assisting in the confocal image acquisition, Tsg-Hui Chang (NINDS) for help with colony management and mice care, Dr. Andrew Griffith (NIDCD) for sharing the anti-TMC1 antibody with us, and Dr. Lisa Cunningham for invaluable comments on the manuscript. This research was supported by the Intramural Research Program of the NINDS, NIH, Bethesda, MD, to K.J.S. and the Mouse Auditory Testing Core at the NIDCD, NIH, Bethesda, MD, to T.S.F. (ZIC DC-000080).

## AUTHOR CONTRIBUTION AND COMPETING INTEREST

**Angela Ballesteros:** Investigation; Conceptualization; Data curation; Formal analysis; Supervision; Validation; Visualization; Writing - original draft and review & editing. **Tracy S. Fitzgerald:** Investigation; Data curation; Formal analysis; Validation; Visualization; Writing - review & editing. **Kenton J. Swartz:** Supervision; Conceptualization; Funding acquisition; Resources; Writing - review & editing.

The authors declare no competing interests.

## SUPPLEMENTARY MATERIAL

**Supplementary figure 1:**
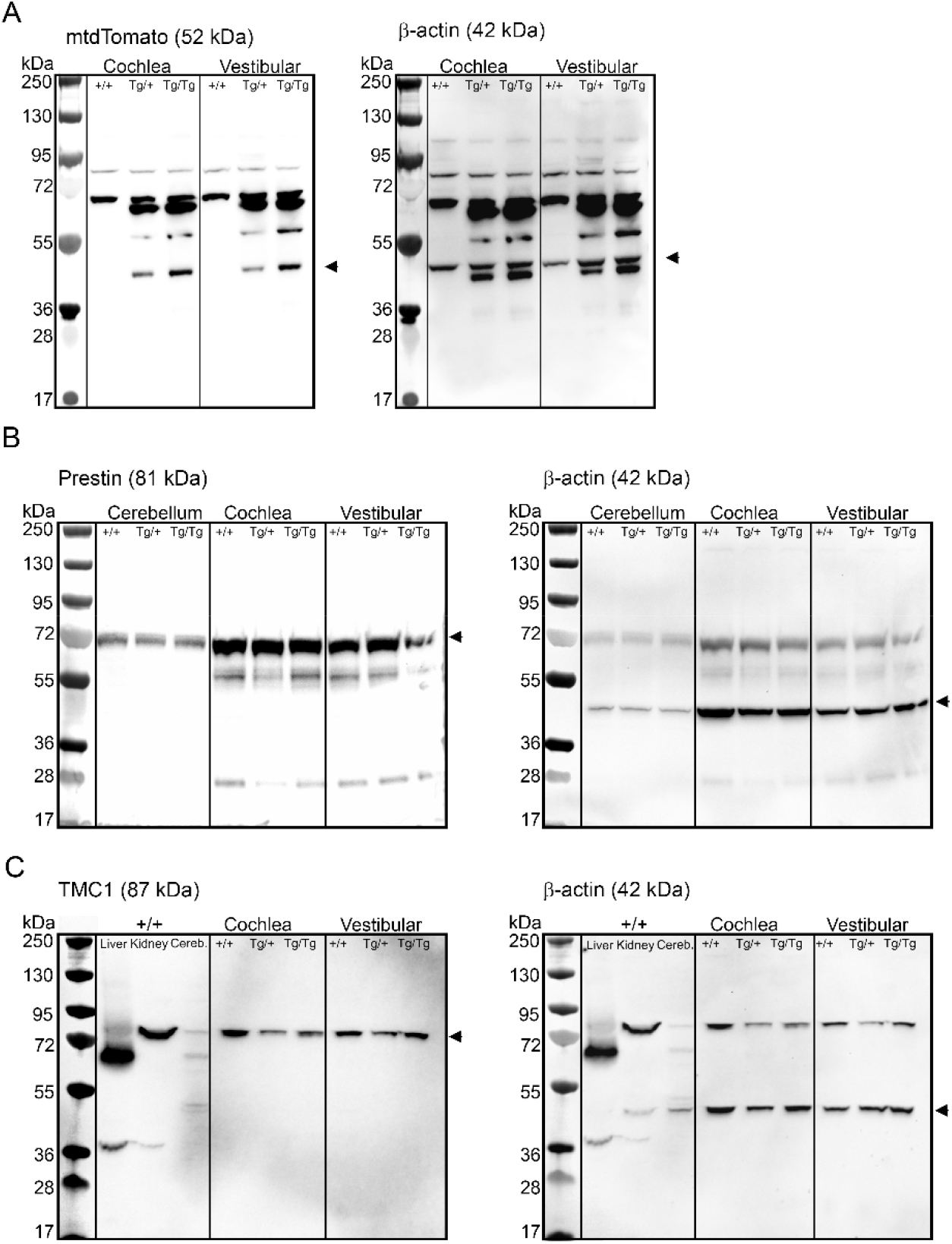
Whole western blot membranes of figures 3D (A), 5A (B), and 7A (C). Western blot for the specific protein (left) and its loading control β-actin (right) are shown. The predicted molecular weight (MW) of the proteins is specified at the top of each membrane, and the band corresponding to the individual protein is indicated with a red arrowhead. Tissue and mice genotype corresponding to the loaded sample is indicated at the top of each lane. Colorimetric image of each membrane showing the protein ladder (MW line) was overlaid with the chemiluminescent western blot for each specific protein for visualization and protein sizing. The MW in kDa of the proteins contained in the ladder is indicated at the left of each blot.

## Notes

### Competing Interest Statement

The authors have declared no competing interest.

